# Urbanization drives genetic erosion and population structure in a historically connected carnivore population

**DOI:** 10.1101/2025.10.03.680202

**Authors:** Laurel E.K. Serieys, Megan Jackson, Shaelynn Sleater-Squires, Gabriella R.M. Leighton, Marine Drouilly, Storme Viljoen, Bogdan Cristescu, Kristine J. Teichman, Deborah J. Winterton, Robert K. Wayne, Jacqueline M. Bishop

## Abstract

Urbanization is a dominant driver of habitat fragmentation globally, creating small, isolated wildlife populations vulnerable to accelerated genetic drift, reduced genetic diversity, and increased population differentiation. We investigated how urban development affects the genetic composition and structure of caracals (*Caracal caracal*) in Cape Town, South Africa using microsatellites and mitochondrial DNA sequence data. Sampling across four geographically disparate urban and rural populations revealed contrasting temporal patterns: mitochondrial markers indicated historical genetic connectivity among populations, while microsatellite data demonstrated recent genetic structuring driven primarily by urbanization. An extensively isolated urban population showed reduced allelic richness and pronounced genetic differentiation, reflecting urbanization as a strong barrier to gene flow. Within the isolated urban population, GPS-collared caracals demonstrated a degree of spatial genetic organization, with related individuals maintaining significantly higher home range overlap despite inhabiting a severely fragmented urban landscape. This kin-structured space use occurred despite caracals in the system having large home ranges compressed within a relatively small, isolated environment. Our findings reveal that urbanization has rapidly disrupted gene flow in this otherwise geographically widespread and adaptable carnivore, imposing a sufficient barrier to generate detectable genetic consequences within contemporary timeframes. The contrasting signals from historical versus contemporary molecular markers highlight urbanization’s role in fragmenting previously connected populations and demonstrates the value of multi-marker approaches for detecting anthropogenic impacts on wildlife populations. These results underscore urbanization’s capacity to rapidly alter population genetic dynamics, even in a highly mobile and adaptable carnivore.

## Introduction

More than 75% of the Earth’s land area has been impacted by the human footprint (Venter et al. 2016), with urbanization, developing at an unprecedented pace globally (Grimm et al. 2008; Seto et al. 2012; Liu et al. 2020), representing a primary threat to biodiversity through habitat loss, degradation, and fragmentation and ultimately population extirpation (McKinney 2002; Grimm et al. 2008; McDonald et al. 2008). When urbanization fragments habitat into patches surrounded by severely altered vegetation, impervious surfaces, and human infrastructure, it reduces vagility among populations (Tucker et al. 2018). These processes create isolated populations that experience reduced gene flow and accelerated genetic drift (Frankham et al. 2002; Frankham 2005; Keyghobadi 2007; Benazzo et al. 2017; Kyriazis et al. 2024). The loss of genetic variation in once-connected populations can reduce the adaptive potential of populations (Keller and Waller 2002; Exposito-Alonso et al. 2022), and, in extreme cases, can lead to inbreeding depression that threatens population persistence (Keller and Waller 2002; Frankham 2005). Genetic diversity is thus crucial to population viability (Benson et al. 2016), especially in the face of environmental change (Frankham 2005; Lourenço et al. 2017; Exposito-Alonso et al. 2022).

Although studies examining the genetic effects of urbanization on wild animal populations remain limited and are primarily focused on North America, research demonstrates that urban development significantly affects genetic structure, diversity, and gene flow across a variety of species and geographies (Miles et al. 2019) . However, responses can vary considerably among taxonomic groups. A continent-wide analysis of North American mammals and birds revealed that mammals in more urban-disturbed habitats exhibited lower effective population sizes and genetic diversity and increased structure, whereas there was no consistent relationship between genetic metrics and habitat disturbances for birds (Schmidt et al. 2020). This finding suggests that terrestrial mammals may be among the taxa most vulnerable to genetic impacts from urban development.

Several mammalian species exemplify genetic sensitivity to urbanization. Increased population genetic structure and reduced genetic diversity associated with urbanization has been documented in white-footed mice (*Peromyscus leucopus*) in New York City, black-tailed prairie dogs (*Cynomys ludovicianus*) in Colorado, Merriam’s kangaroo rats (*Dipodomys merriami*) in New Mexico, and gray squirrels (*Sciurus griseus*) in California (Magle et al. 2010; Munshi South et al. 2016; Hurtado and Mabry 2019; DeMarco et al. 2021). Given their large space requirements and low population densities, medium-to large-sized carnivores can be especially sensitive to the ecological and genetic effects of habitat fragmentation via urbanization (Crooks 2002; Murphy et al. 2017). In Los Angeles, urban bobcat (*Lynx rufus*) and coyote (*Canis latrans*) populations separated by a major freeway experienced less gene flow than expected based on the number of individuals recorded crossing the freeway, possibly due to home range pile-up against hard urban infrastructure reducing the ability of those that crossed the freeway to successfully establish territories and find mates (Riley et al. 2006). The genetic consequences of habitat fragmentation and population isolation on carnivores are known to include reduced genetic diversity, inbreeding, and enhanced population structure (Serieys et al. 2015; Benson et al. 2019; Gustafson et al. 2019; Adducci et al. 2020; Exposito-Alonso et al. 2022) ultimately threatening some populations with reduced fitness and local extinction (Benson et al. 2016, 2019). Overall, while genetic responses to urbanization range from significant population divergence and genetic erosion to negligible changes depending on taxa, terrestrial mammals, and carnivores in particular, appear to be particularly susceptible to urban-induced genetic changes (Benson et al. 2016, 2019; Johnson and Munshi-South 2017; Miles et al. 2019).

Caracals (*Caracal caracal*) are a medium-sized felid that has one of the most widespread geographic distributions of any wild felid, illustrating their adaptability to a variety of ecosystems (Hunter 2015). They have large home ranges and as solitary carnivores have low population densities (Sunquist and Sunquist 2002; Hunter 2015) making them vulnerable to the genetic effects of habitat fragmentation (Murphy et al. 2017; Kyriazis et al. 2024; Gargiulo et al. 2025). Despite their widespread geographic distribution ranging from Africa to Asia, few studies have used genetic markers to investigate caracal demographic patterns, genetic diversity, or population structure across fragmented landscapes. In South Africa, caracals are distributed throughout the country. To our knowledge, three genetic studies have been conducted on the species in the country, examining the genetic effects of lethal management, local-scale population connectivity, and isolation by urbanization (Tensen et al. 2018, 2019; Kyriazis et al. 2024). In one study using genomic sequencing, Kyriazis et al. (2024) found limited gene flow into an urban caracal population isolated by the city of Cape Town. Low levels of migration into the urban population were documented with an estimated rate of 1.3 effective migrants per generation over a period of approximately 75 years. The study also found an effective population size of only 28 individuals. Not surprisingly, inbreeding was evident as long runs of homozygosity (Kyriazis et al. 2024). This study points to the powerful genetic effects of isolation by urbanization even over relatively short timescales that could with time lead to inbreeding depression. Other threats to this population include vehicle collisions, poaching, lethal management, domestic dog attacks and disease and pesticide exposure (Serieys et al. 2019; Viljoen et al. 2020; Leighton et al. 2022a; Serieys and Leighton, unpubl.data).

Here, we use microsatellite and mitochondrial markers to assess the influence of isolation by urbanization on the contemporary and historical population genetics of caracal in Cape Town, South Africa, building on the study by Kyriazis et al. (2024). Given the complexity and varying impacts of urban disturbance on genetic metrics across species, it is essential to utilize informative genetic tools to unravel changes in the genetic composition and connectivity of populations. Microsatellites and mitochondrial sequences are powerful genetic markers that can offer complementary insights into the population genetic effects of human activities (Wayne and Morin 2004). We use samples collected from an isolated urban population, as well as three contiguous outgroup populations in western and central South Africa, to evaluate contemporary and historical population structure and genetic diversity metrics. Using microsatellite makers on a subset of individuals that were GPS-collared in the urban, isolated population, we also take the opportunity to test whether individuals that have higher pairwise relatedness values also exhibit greater home range overlap, hypothesizing that because the caracals inhabit a relatively small and isolated area that we will not see greater home range overlap amongst individuals with higher relatedness. We investigate how contemporary (microsatellite) population structure across the four populations compares with historical (mitochondrial) patterns of population structure.

By integrating data from two types of genetic markers, we gain a comprehensive understanding of how urbanization influences recent genetic patterns while gaining insight into historical patterns of population structure. This approach elucidates evolutionary pressures on a solitary carnivore in a fragmented and isolated habitat and establishes a framework for future urban wildlife research. Our study also contributes to the limited body of work that has examined the impacts of urbanization on the genetics of natural wildlife populations in Africa (Fusco et al. 2021).

## Methods

### Study areas

Caracals were sampled from four regions (Figure 1): 1) the Cape Peninsula (CP), 2) the Greater Cape Town (GCT) area, 3) the Central Karoo (CK), and 4) Namaqualand (NMQ). The Cape Peninsula encompasses 320 km^2^ of available wildlife habitat and is isolated by the dense urban Cape Town matrix spanning roughly 800 km^2^ (Figures 1-2). The Cape Peninsula is primarily comprised of Table Mountain National Park (TMNP; 250 km^2^) and city of Cape Town reserves within the Cape Peninsula. Development within the Cape Peninsula primarily includes residential and commercial areas and vineyards. We sampled caracals across the Greater Cape Town region, which included the False Bay Nature Reserve, a small nature reserve comprising <3km^2^ within Cape Town’s urban matrix and surrounded by the densely populated region of Cape Town (9,000–17,000 people/km^2^). The urban matrix that now isolates the Cape Peninsula has undergone development and expansion since the 1850s, starting in the northern most section of the Cape Peninsula (Figure 2). Expansion of the urban matrix has continued since then with a large swath of urban matrix developed by 1977 (see Figure 2 for the progression of development, City of Cape Town, 2023). Additional development in the 1980s through roughly 2010 effectively isolated the Cape Peninsula from larger contiguous areas inhabited by caracals.

**Figure 1.**
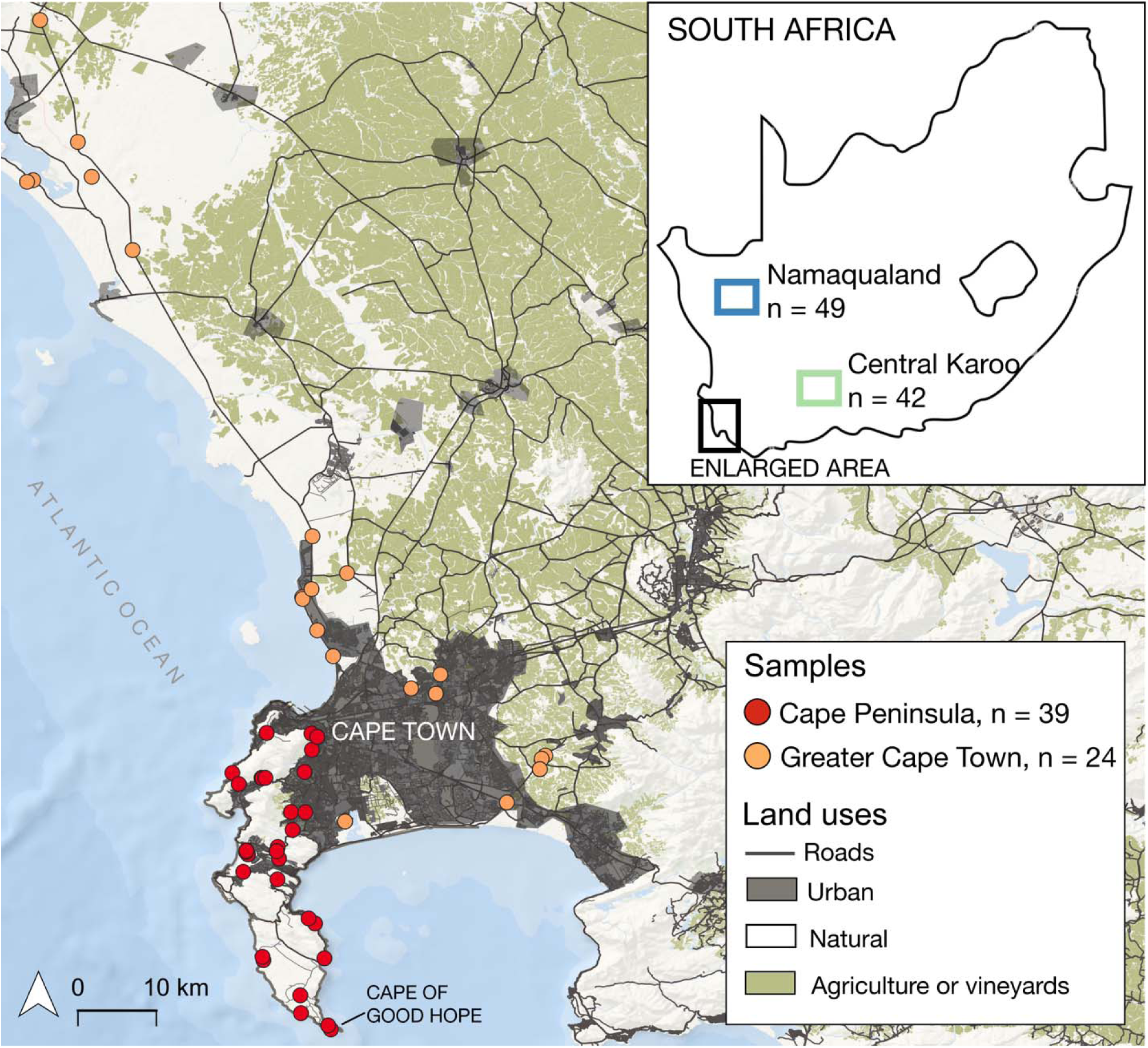
Map of the study areas in South Africa showing sampling locations of caracals (Caracal caracal). Sample sizes represent microsatellite genotypes.

**Figure 2.**
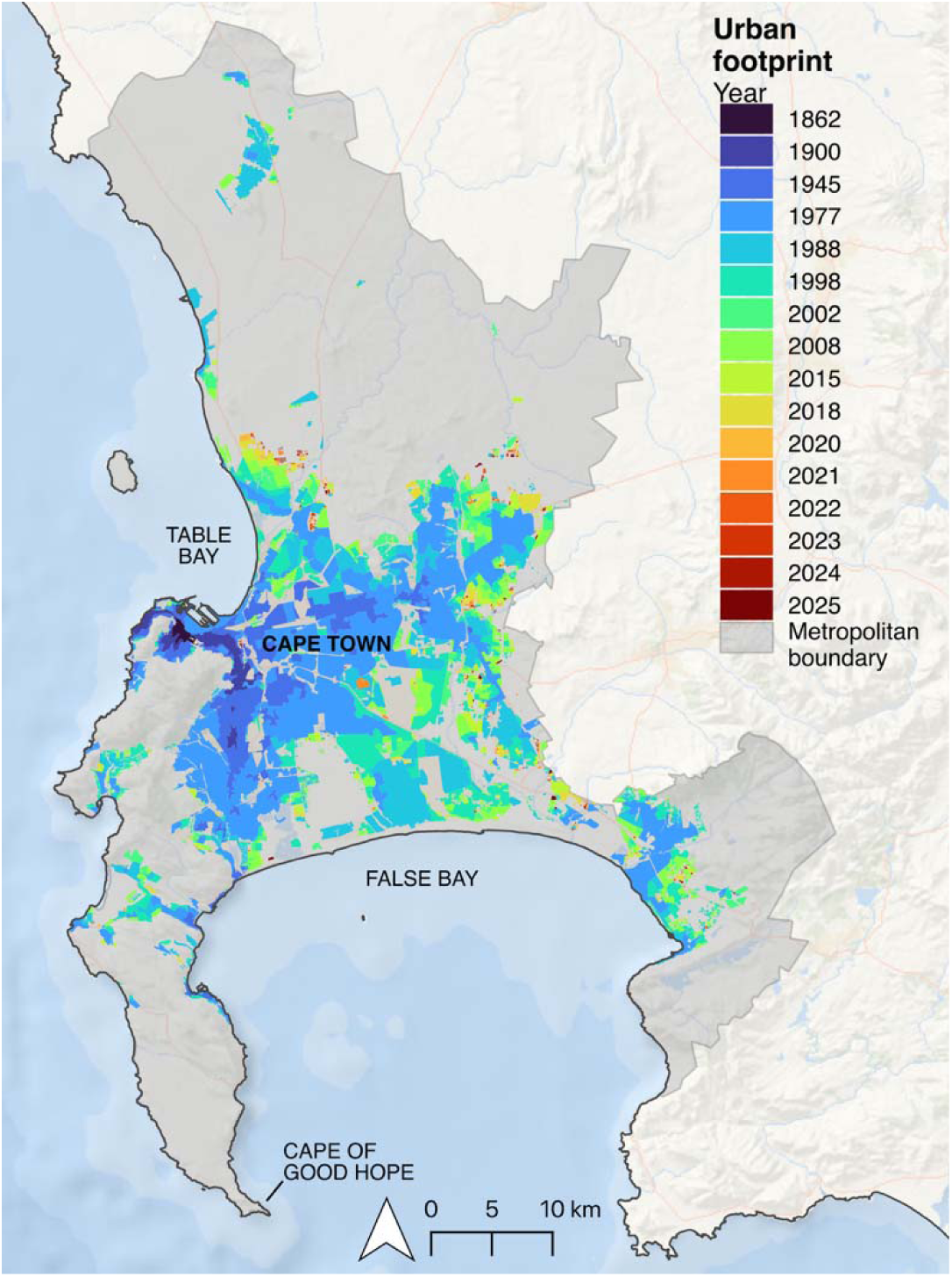
Map of the development of the urban footprint of Cape Town since 1862 showing the progression of isolation of the Cape Peninsula. Map modified from City of Cape Town, 2023 and Cape Town’s urban footprint dataset were used to create the map.

Overall, for the purposes of our study, the Greater Cape Town region largely comprises dense Cape Town urbanization, suburban development, agricultural areas, vineyards, city nature reserves, and West Coast National Park, roughly 90 km north of Cape Town (Figure 1). Natural areas in both the CP and GCT are dominated by the fynbos biome which is a low-growing, dense shrubland (Rebelo et al. 2011).

The Central Karoo is a rural region located approximately 350 km north-east of Cape Town, while Namaqualand is a rural region located approximately 480 km north of Cape Town (Figure 1). Both regions are semi-desert areas that consist mainly of privately-owned small-stock farmland. Human population densities are low (CK: 1.8 people/km^2^; NMQ: 1.1 people/km^2^) and croplands are somewhat rare and are often restricted to small areas immediately adjacent to farmhouses. Livestock primarily comprises free-ranging sheep and goats that feed on native vegetation. While overall dominated by livestock farmland, the Namaqualand area includes the protected Namaqua National Park. Both the Central Karoo and Namaqualand study area landscapes are contiguous, unlike the Cape Peninsula and greater Cape Town area.

### Sampling

All samples (blood, skin, muscle, or liver tissue) were collected between 2014 – 2022. Between 2014 – 2017, we intensively trapped caracals using cage traps for spatial ecology studies across the Cape Peninsula and within the False Bay Nature Reserve in the Greater Cape Town region (Leighton et al. 2022b; Serieys et al. 2023). Caracals were immobilized as in Serieys et al. (2023) and a veterinarian assisted at all captures. While caracals were immobilized, we collected blood and a small (3mm) piece of ear tissue. Within both the Cape Peninsula and the Greater Cape Town region, we additionally collected samples opportunistically from mortalities (primarily vehicle mortalities) during 2015 – 2025 from which we collected blood (when possible), skin, muscle, and liver tissue. In the Central Karoo, muscle and skin tissue samples were collected as in Tensen et al. 2018 and Viljoen et al. 2020 from animals culled by local farmers and professional hunters during standard permitted culling operations. In Namaqualand, we also live-trapped caracals from 2014 – 2015 using cage traps and padded foothold traps as in Teichman et al. 2023. Blood samples used in this study were collected from Namaqualand caracals during captures. In addition, Namaqualand skin tissue samples were collected from local farmers that had pelts in their possession from permitted hunting operations. No animals were killed for the purpose of this study.

### Ethics Statement

All caracal trapping, handling, and sampling protocols followed ethical guidelines approved by the University of Cape Town (2014/V20/LS), Cape Nature (AAA007-00147-0056 [L.Serieys], AAA007-00560-00161 [S.Viljoen], AAA007-00074-0056 [M.Drouilly]), and CN44-87-33288 [G.Leighton], and South African National Parks (SANParks) (SERL/AGR/017– 2014/V1). Additionally, for Namaqualand, capture and handling protocols were conducted with permits from the Northern Cape Department of Environment and Nature Conservation (permit 1157/2013), SANParks (permit CRC-2013/029-2014), the University of British Columbia Okanagan (A15-0204), the University of Cape Town (2013/V30/BC), and Stellenbosch University (SU-ACUM14-00001). No permits were required to sample pelts in Namaqualand.

### DNA extractions and microsatellite genotyping

DNA was extracted from blood, skin, muscle or liver tissue using the QIamp® DNA Mini Kits (Qiagen, Germantown, MD, USA) according to the manufacturer’s protocols. Individuals were genotyped using 15 supposed neutral microsatellite markers including dinucleotide markers (FAC008, FCA023, FCA031, FCA043, FCA045, FCA077, FCA082, FCA090, FCA096, FCA126 and FCA132), a trinucleotide marker (FCA741), and tetranucleotide markers (FCA391, FCA742, and BC1AT). Each of these markers was developed for the domestic cat (*Felis catus*, Menotti-Raymond et al. 1999, 2003)), except for BC1AT, which was developed for the bobcat (Faircloth et al. 2005). Microsatellite genotyping was performed using polymerase chain reaction (PCR) amplification methodologies adapted from Boutin-Ganache et al. 2001 with the QIAGEN Multiplex PCR Kits (QIAGEN, Valencia, CA, USA). No changes were made to the reverse primers, but the forward primers were modified to contain the 16-bp M13 universal sequence (−20: 5’-GTC AAA CGA CGG CCA G-3’) at the 5’ end for all loci. A second primer, a dye-label (6-FAM or NED, Applied Biosystems, Waltham, MA, USA) attached to the M13, was also added to the PCR. Each amplification reaction was made up of 1.0µl M13 hybrid primer mix, 2.1µl double distilled H_2_0, 0.4µl 10mg/ml bovine serum albumin, 5.0µl QIAGEN multiplex PCR master mix and 1.5µl (20 – 50ng) DNA, to a final reaction volume of 10µl. PCR reactions were performed using the following thermocycling profile: 15min at 95°C for the initial activation step, followed by 25 cycles of 30 s at 94°C, 90s at 59°C and 60s at 72°C. This was followed by 15 cycles of 30 s at 94°C, 90 s at 53°C and 60 s at 72°C and the final step was 30 min at 60°C. PCR products were sized by running a 1:20 dilution with 9.7 ul Hi-Di formamide and 0.3 ul GeneScanTM 500 LIZ® Size Standard 10 on a capillary 3730 DNA Analyzer (Applied Biosystems, Waltham, MA, USA). PEAK SCANNER 2.0 software (Applied Biosystems, Waltham, MA, USA) was used to score allele sizes. Samples were re-genotyped if they had ambiguous allele calls or if amplifications failed. Positive and negative controls were used to ensure consistent genotyping calling.

### Validating and characterizing microsatellite data

We tested each locus for deviation from Hardy-Weinberg equilibrium (HWE) and linkage disequilibrium (LD) using GENEPOP v4.7.5 (Rousset 2008). We tested each marker pair for LD both within populations, and across all populations. We made >100 comparisons and thus we used the Bonferroni method to correct the critical value corresponding to α = 0.05 for 105 comparisons (α = 0.0005) (Rice 1989). We conducted global tests of heterozygote deficiency and excess for each population and for each locus, and because 30 tests were conducted per population, we corrected for multiple tests (α = 0.0017). We evaluated genotyping error, the presence of null alleles, scoring errors, and allele dropout using MICRO-CHECKER (Oosterhout et al. 2004).

### Microsatellite genetic diversity and relatedness

We calculated genetic diversity measures including observed (H_O_) and expected (H_E_) heterozygosities (Nei 1987), per locus allelic richness (A_R_), and the inbreeding coefficient (F_IS_) using the R package ‘hierfstat’ (Goudet 2005). We quantified individual pairwise relatedness for each population using Maximum-likelihood (ML)-RELATE (Kalinowski et al. 2006).

To test for significant differences for each value across populations, we first tested for normality for each value using a Shapiro-Wilk test. Next, we performed a Kruskal-Willis or an ANOVA for each value across populations. Finally, pairwise comparisons were calculated with Wilcoxon rank sum or t-tests (α = 0.05). All analyses were performed in R v.4.3.2 (R Development Core Team, 2023).

### Population structure and genetic differentiation

We assessed population structure using a multivariate analytical approach and a Bayesian clustering method. First, we performed a discriminant analysis of principle components (DAPC), a multivariate approach that identifies and describes clusters of genetically related individuals (Jombart et al. 2010) using ‘adegenet’ (Jombart 2008) in R (R Development Core Team, 2023). We used the method to test for genetic differences among *a priori* defined populations (k = 4), starting with an input of 60 principal components (chosen according to the plot of variance explained by the PCA) and three discriminant functions (k – 1). We performed cross validation to assess correct assignment rates.

We next implemented STRUCTURE 2.3.4 (Pritchard et al. 2000) which can infer population assignments without *a priori* assumptions about sample location. Using STRUCTURE, we first inferred the number of genetic clusters (K) without information about sampling locations. We evaluated the stability of inferred clusters using 10 independent runs (K = 1–10) with a burn in period of 50,000 iterations and 500,000 MCMC cycles. Next, using STRUCTURE HARVESTER (Earl and vonHoldt 2012), we examined raw probability values of LnP(K) and the ΔK estimate (Evanno et al. 2005) to help evaluate the optimal number of genetic clusters. The most likely population clustering agreed with our geographic sampling locations, and thus we performed the analyses again but with sampling location as prior information to evaluate the individual cluster assignments. We report these results in Figure 3. To align the multiple STRUCTURE replicates, we used CLUMPP 1.1 (Jakobsson and Rosenberg 2007). Last, we used STRUCTURE to identify potential migrants, individuals that are assigned to a population different from the one in which they were sampled, between the adjacent Cape Peninsula and Greater Cape Town area (Berry et al. 2004). Individuals were considered migrants if their assignment and posterior probabilities were >50% to a population different than the one in which they were sampled.

**Figure 3.**
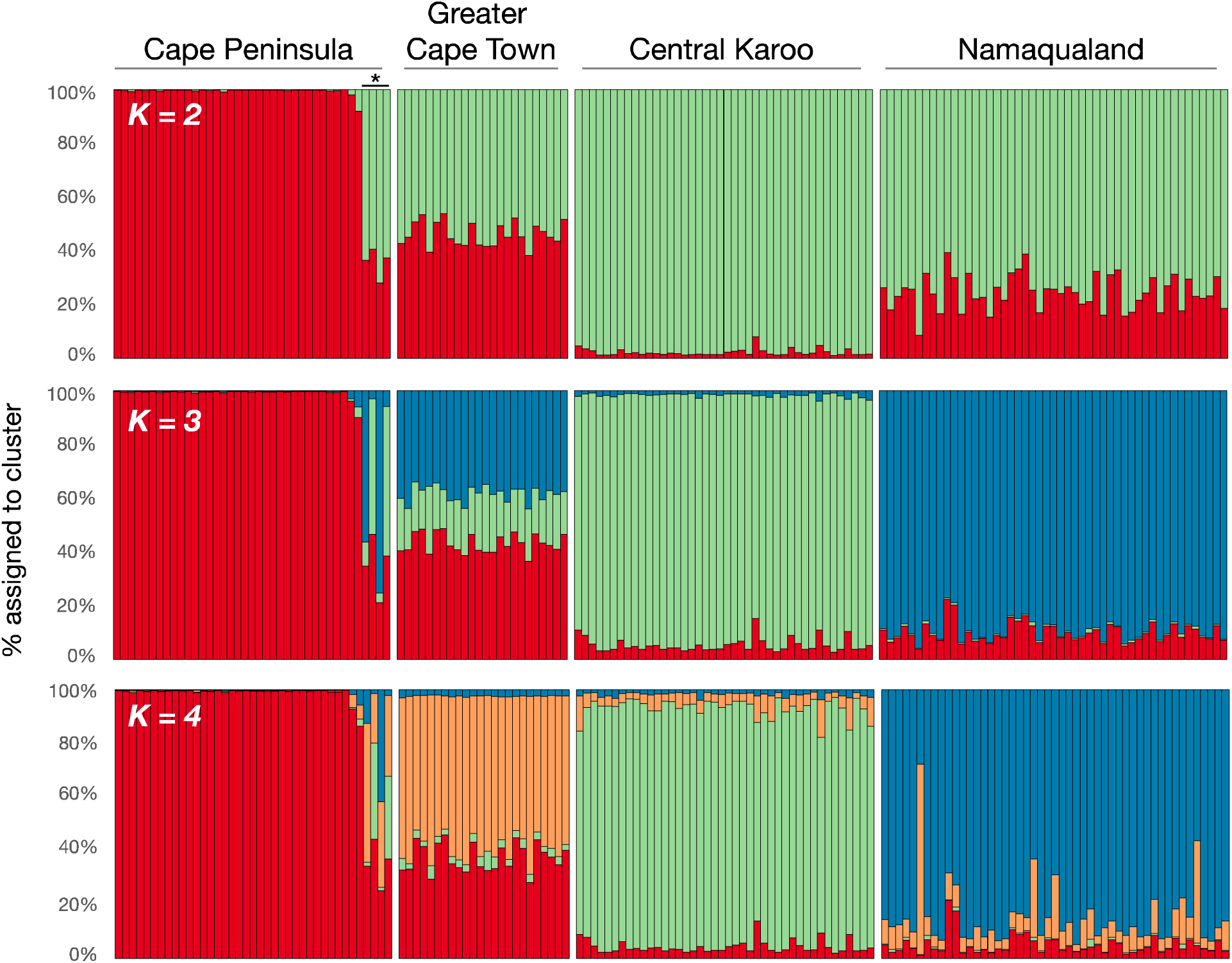
Results of the population structure analysis, using capture location as prior information, using microsatellite loci. Each vertical bar represents a single individual and the shading of each bar corresponds to the probability of genetic assignment to one of four populations of caracals. Asterisk above four individuals in the Cape Peninsula represent first- or second-generation migrants. While the Evanno method to choose the most likely number of population clusters based on ΔK identified K=2 as the optimal K-value, K=4 aligns best with our knowledge of the geographic location of populations.

We performed analysis of molecular variance (AMOVA) to assess structuring within the data. We also quantified genetic differentiation between populations using D_est_ (Jost 2008), R_ST_ (Slatkin 1995), and F_ST_ as calculated using the AMOVA. D_est_ is a frequency-based method that accounts for differences in allelic diversity for highly polymorphic microsatellite markers (Jost 2008) while R_ST_ (Slatkin 1995) is a measure, inferred from allele frequencies, of the extent of population differentiation that takes explicit account of the mutation process at microsatellite loci. All three values were calculated in GenAlEx 6.51 (Peakall and Smouse 2006, 2012). D_est_ was calculated with 999 bootstraps, while probabilities for both D_est_ and F_ST_ were computed using 999 permutations to test for significance.

### Home range overlap and relatedness

We tested whether GPS-collared individuals in the Cape Peninsula that had higher relatedness values also had greater home range overlap. We estimated 95% LoCoH-a home ranges implemented in the R package ‘t-locoh’ (Lyons et al. 2013) as described in more detail in (Leighton et al. 2022b). Next, we performed a Pearson correlation, in R (R Core Team, 2023), to test whether pairwise relatedness correlated with average pairwise 95% LoCoH-a overlap values.

### mtDNA control region sequencing and data analysis

To explore signatures of historic connectivity across the study area we also sequenced a section of the mitochondrial (mtDNA) control region for individuals from each study area using primers LO forward 5’ – CCCAAAGCTGAAATTCTACTTAAA CTA – 3’ and E3 reverse 5’ – ATGACCCTGAAGAAASAACCAG – 3’ (Smit et al. 2008). PCR amplification was performed using DreamTaq Green PCR Master Mix (Thermo Fisher Scientific, Waltham, MA, USA) in a final reaction volume of 20 µl containing 1 µl (0.5 μM) of each primer, 10 µl DreamTaq Green PCR Master Mix; 7 µl of double distilled H_2_O, and 1-3 µl (∼10ng) of template DNA. PCR reactions were performed on a GeneAmp 9700 thermocycler (Applied Biosystems, Waltham, MA, USA) using the following protocol: 3 min at 95°C, 35 cycles of 95°C for 30s, 54°C for 30s, and 72°C for 30s, and 72°C for 10 min. PCR products were sequenced and analyzed in both directions at the Central Analytical Facility, Stellenbosch University using BigDye 3.1 chemistry on an ABI 3730xl (Applied Biosystems, Waltham, MA, USA) Genetic Analyzer. Sequence fragments were trimmed to 741 base pairs of comparable sequence data for further analysis.

Sequences are available on GenBank (accession numbers XXX-XXX). Control region DNA electropherograms were scored and then aligned in Geneious Prime 2025.2.1 (www.geneious.com) using both the Geneious and ClustalW alignment algorithms. Summary diversity statistics for haplotype and nucleotide diversity were calculated in DnaSP v6.12 (Rozas et al. 2017). We used haplotype network analysis to explore genealogical relationships and spatial distribution of control region haplotypes using the NeighborNet splits method (Bryant and Moulton 2004) in SplitsTree4 (Huson and Bryant 2006, 2024). An unrooted NeighborNet network was constructed using the full alignment of haplotypes identified in the study; the analysis used uncorrected ‘p’ distances and network splits, and the resulting structure was tested with 1000 bootstrap replicates

## Results

### Microsatellite caracal sampling

We obtained tissue samples from 154 caracals from four regions (CP, n = 39; GCT, n = 24; CK, n = 42; NMQ, n = 49). Of the 154 individuals that were genotyped using 15 microsatellite loci, 31 were missing data for one locus, 10 were missing data for two loci, three were missing data for three loci, and one individual was missing data for four loci (Supplementary Table S1).

No genotyping error or allele dropout was observed. Six loci showed evidence of null alleles, although no loci showed null alleles across all four populations (Supplementary Table S2). However, six loci (CP: FCA045 and FCA090; GCT: FCA096; CK: FCA045 and FCA090; NMQ: FCA031, FCA090, and FCA132; Supplementary Table S2) showed high frequencies of null alleles (>0.10) in individual populations. While the high estimates of null alleles observed could lead to overestimated genetic differentiation between populations, true null alleles are generally only found in populations exceeding effective population sizes of 50,000. Further, if the genotyped portion of the population is not thoroughly representative of the focal population, the frequency of null alleles can be artificially inflated (Chapuis and Estoup 2007).

Five of 420 pairs of loci demonstrated significant LD after correction for multiple tests (Supplementary Table S3). Yet none of these pairs were in LD across all four populations, suggesting the significant LD values were likely the result of population structure, particularly because four of the five pairs in LD were observed in the Cape Peninsula population. Two of the 15 loci deviated from HWE, although none deviated from equilibrium across all four populations. We observed significant heterozygote deficiencies in FCA090, FCA045, and FCA391, although not across all four populations (Supplementary Table S4). We did not observe heterozygote excess for any locus for any population.

### Microsatellite genetic diversity

All loci were polymorphic with 5 – 13 alleles observed per locus. Private alleles were observed in all populations (CP, n = 1; GCT, n = 2; CK, n = 9; NMQ, n = 29). Mean per-locus allelic richness (A_R_) ranged from 3.52 in the Cape Peninsula population to 5.77 in the Namaqualand population (Table 1) and differed significantly across the four populations (X^2^= 16.94, p < 0.001). Allelic richness was significantly lower in the Cape Peninsula when compared with Greater Cape Town (t = −2.179, p = 0.038), Central Karoo (W = 33.0, p = 0.001), and Namaqualand (t = −4.2104, p <0.001). Observed heterozygosity (H_O_) ranged from 0.50 – 0.64 but these values did not significantly differ across the four populations (W = 4.84, p = 0.184). In contrast, expected heterozygosity (H_E_) ranged from 0.51 – 0.69 (Table 1) with significant differences across some populations (W = 10.13, p = 0.017). The Cape Peninsula population had lower H_E_ than the Greater Cape Town (t = −2.13, p = 0.043), the Central Karoo (W = 44.5, p = 0.005), and Namaqualand (t = −2.56, p = 0.016) populations, but there were not significant differences between the other populations. The inbreeding coefficient (F_IS_) ranged from 0.06 (CP and GCT) to 0.10 (CK and NMQ) (Table 1) and did not significantly differ across the populations (W = 1.07, p = 0.784).

**Table 1.** Mean genetic diversity measures: allelic richness (AR), observed heterozygosity (H_o_), expected heterozygosity (H_e_), and relatedness (R) for each population. Standard error values are in parentheses.

| Population | N | AR | $H_o$ | $H_e$ | $F_{IS}$ | R |
| --- | --- | --- | --- | --- | --- | --- |
| Cape Peninsula | 39 | 3.52 (0.32) | 0.50 (0.06) | 0.51 (0.04) | 0.06 (0.07) | 0.12 (0.005) |
| Greater Cape Town | 24 | 4.63 (0.39) | 0.60 (0.04) | 0.63 (0.03) | 0.06 (0.04) | 0.06 (0.005) |
| Central Karoo | 42 | 5.37 (0.46) | 0.64 (0.05) | 0.69 (0.03) | 0.10 (0.05) | 0.06 (0.003) |
| Namaqualand | 49 | 5.77 (0.43) | 0.61 (0.05) | 0.65 (0.03) | 0.10 (0.05) | 0.06 (0.002) |

Relatedness differed significantly across populations (W = 38.02, p <0.001). In the Cape Peninsula, relatedness was twice as high (R = 0.123, SE = 0.005) as the Greater Cape Town (R = 0.058, SE = 0.005), Central Karoo (R = 0.062, SE = 0.003), and Namaqualand (R = 0.061, SE = 0.002) populations (Table 1). Pairwise comparisons of relatedness revealed a significant difference between the Cape Peninsula and all other populations (all pairwise comparisons p <0.001), but also between the Greater Cape Town and Namaqualand (p = 0.031) populations. However, the significance of the latter likely reflects the high number of individual pairwise relatedness observations despite the similarity of the Greater Cape Town and Namaqualand relatedness values.

### Population structure and genetic differentiation

The Bayesian clustering method implemented in STRUCTURE indicated that two genetic clusters were most likely using the ΔK approach. The two genetic clusters (K = 2) aligned with the Cape Peninsula forming one cluster, and the other three regions forming the second cluster (Figure 3). Importantly, although the Cape Peninsula and the Greater Cape Town area are adjacent to each other (Figure 1), separated by approximately 3 km, the Greater Cape

Town area fell into a different genetic cluster than the Cape Peninsula. However, population clusters resolved at K = 4 corresponded better with the geographic distribution of the populations (e.g., Namaqualand and Central Karoo > 300 km from the Cape Peninsula and Greater Cape Town areas; Figure 3). Further, Evanno et al. (2005) advise that ΔK can be used to help identify the correct number of genetic clusters, but that it should not be the sole method for cluster delineation. DAPC clustering supported four clusters in the dataset that corresponded geographically (Figure 4). The DAPC posterior assignment probabilities of individuals to the four geographically defined clusters was high (81% successful assignment rate) providing additional support for four geographically distinguished populations in the dataset. Using STRUCTURE, we detected four genetic migrants from the Greater Cape Town population into the Cape Peninsula population (Figure 3).

**Figure 4.**
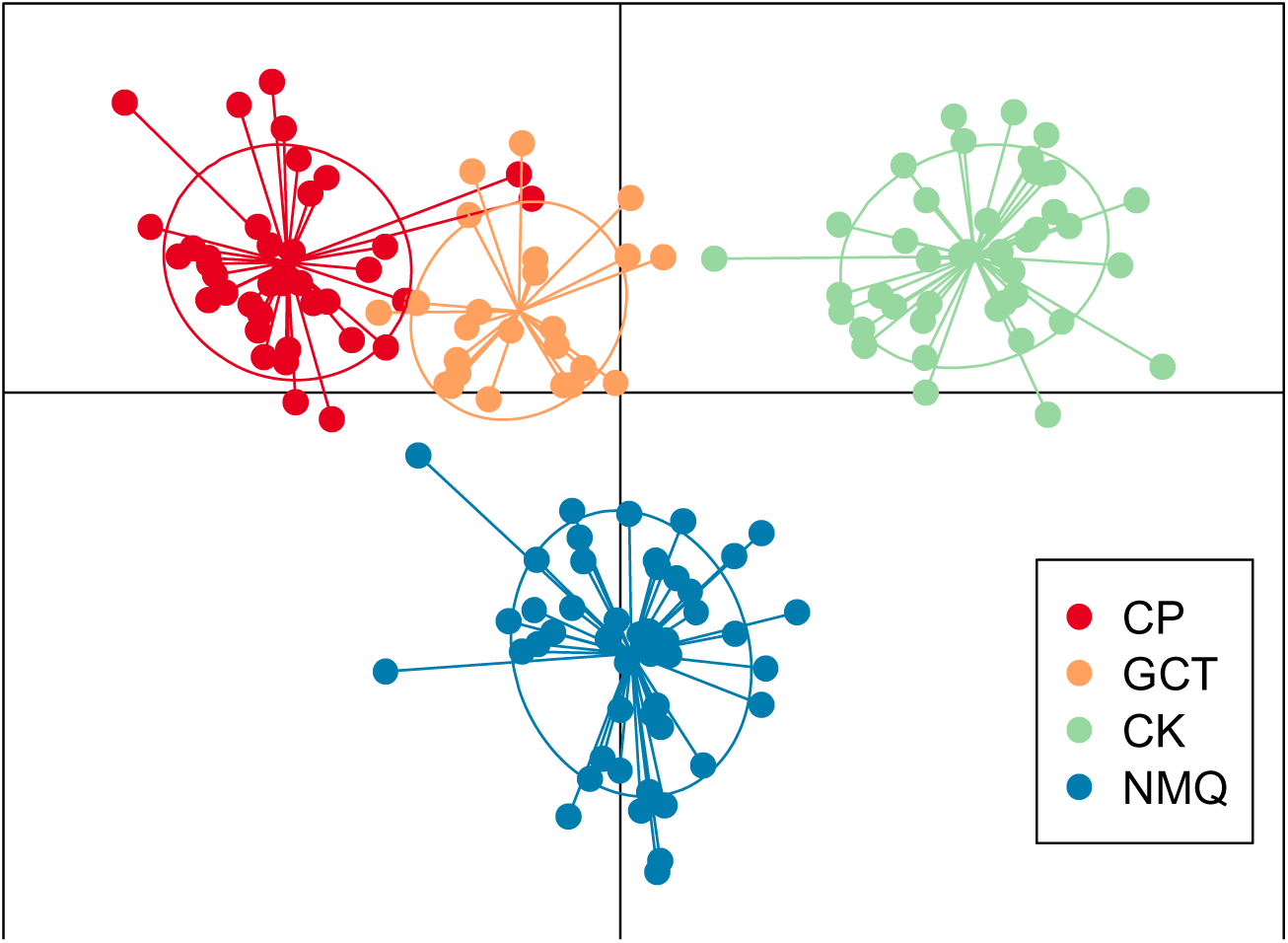
DAPC plot based on microsatellite genotypes showing evident population structure among the four sampled regions. The DAPC posterior assignment probabilities of individuals to the four geographically defined clusters was high (81% successful assignment rate).

For the microsatellite data, the AMOVA supported the geographical structuring of the populations (F_ST_ = 0.08, p = 0.001) with 8% among population variation. Significant genetic differentiation between populations was evident using multiple measures. D_est_ ranged between 0.08–0.27, R_ST_ ranged between 0.05–0.15, and F_ST_, as calculated using an AMOVA, ranged between 0.04–0.14 (Table 2). The trends across each measure were generally consistent. The greatest degree of genetic differentiation was between the Cape Peninsula and Central Karoo populations while the lowest was between the Greater Cape Town and Namaqualand populations.

**Table 2.** Genetic differentiation values across three measures. Probability values are in parentheses. F_ST_ values were calculated using an AMOVA. CP: Cape Peninsula, GCT: Greater Cape Town, CK: Central Karoo, NMQ: Namaqualand.

| Population pair | $D_{est}$ | $R_{ST}$ | $F_{ST}$ |
| --- | --- | --- | --- |
| CP – GCT | 0.09 (0.001) | 0.05 | 0.06 (0.001) |
| CP – CK | 0.27 (0.001) | 0.14 | 0.14 (0.001) |
| CP – NMQ | 0.16 (0.001) | 0.09 | 0.10 (0.001) |
| GCT – CK | 0.15 (0.001) | 0.15 | 0.07 (0.001) |
| GCT – NMQ | 0.08 (0.001) | 0.07 | 0.04 (0.001) |
| CK – NMQ | 0.13 (0.001) | 0.05 | 0.05 (0.001) |

### Home range overlap and relatedness

We estimated 95% LoCoH home ranges for 26 individuals in the Cape Peninsula and found that the proportion of home range overlap ranged from 0–1 while relatedness ranged from 0.00 – 0.86. We found a positive correlation (*r* = 0.104) between pairwise relatedness and average pairwise home range overlap (t = 2.36, p = 0.012) indicating that those with higher pairwise relatedness also had higher average home range overlap.

### mtDNA

We sequenced a total of 113 individuals across our study areas (CP, n = 30; GCT, n = 31; CK, n = 31; NMQ, n = 21) and found 48 haplotypes (overall haplotype diversity H_D_ = 0.88; SD ±0.02). Sequences were characterized by a combination of nucleotide polymorphisms, insertion-deletion mutations, and 47 of 113 (41.6 %) caracals sequenced contained an 80-bp duplication, also reported in Tensen et al. (2018). When compared across our study areas, haplotype diversity was equivocal to overall H_D_ and ranged from H_D_ = 0.73 – 0.89 (SD ±0.02 – 0.08), while nucleotide diversity ranged from π = 0.0132 – 0.0596 (SD ±0.003 – 0.008), and π = 0.0486 (SD ±0.003) for the overall dataset (Table 3). Network analysis did not support any geographic structure in the dataset but rather identified two clades comprising haplotypes from all four study areas (Figure 5). While some haplotypes were specific to a sampling region (CP=7; GCT=7; CK=13; NMQ=15), many were shared across two or more areas. While our focal study area of the Cape Peninsula had seven unique haplotypes out of a total of 11 (63.6 %), its shared haplotypes were common to combinations of the other three study areas.

**Figure 5.**
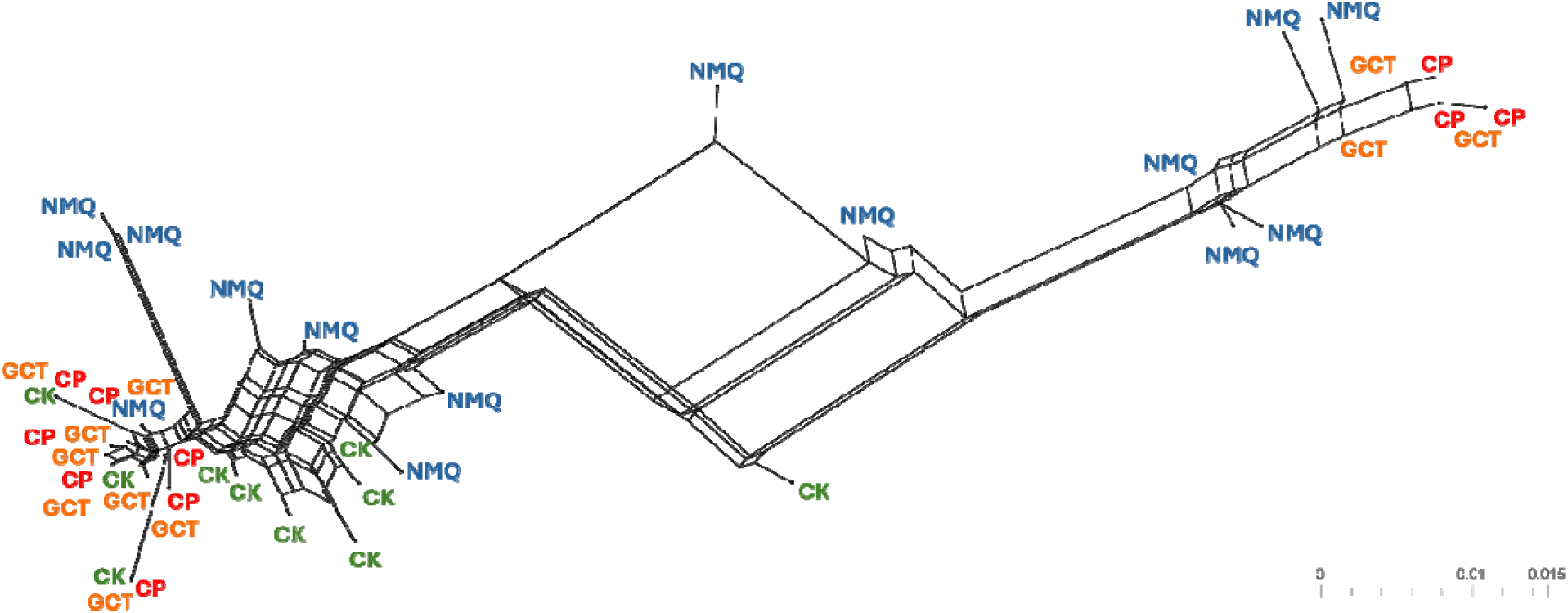
Mitochondrial DNA network analysis figure. Network analysis did not support any geographic structure in the dataset. Instead, the analysis identified two clades comprising haplotypes for all four study areas.

**Table 3.** Mitochondrial control region diversity for caracals (Caracal caracal) from South Africa. H = number of haplotypes; H_D_ = haplotype diversity; π = nucleotide diversity; results provided ± SD. H (number of haplotypes) is not expected to total 48 because some haplotypes are shared across the study areas.

| Study area | No. of samples | H | $H_D$ | $\pi$ |
| --- | --- | --- | --- | --- |
| Cape Peninsula | 30 | 11 | $0.81 \pm 0.03$ | $0.0574 \pm 0.008$ |
| Greater Cape Town | 31 | 13 | $0.88 \pm 0.03$ | $0.0167 \pm 0.003$ |
| Central Karoo | 31 | 13 | $0.73 \pm 0.08$ | $0.0132 \pm 0.003$ |
| Namaqualand | 21 | 18 | $0.89 \pm 0.02$ | $0.0596 \pm 0.004$ |
| All samples | 113 | 48 | $0.88 \pm 0.02$ | $0.0486 \pm 0.003$ |

## Discussion

The accelerating pace of global urbanization presents substantial obstacles to maintaining biodiversity within urban ecosystems. Urban development fundamentally alters evolutionary processes that shape genetic diversity, as fragmented and isolated habitats modify patterns of genetic drift, gene flow, and natural selection (Miles et al. 2019; Fusco et al. 2021). Here we investigated patterns of both historical and contemporary population structure and the effects of genetic drift in a highly adaptable generalist carnivore. Comparing historical and contemporary markers offers insights into how populations have changed over time, including the impact of events like habitat fragmentation, urbanization, and isolation. While mitochondrial DNA analysis reflects extensive historical connectivity among all four study areas, microsatellite data demonstrate pronounced contemporary population structure, particularly isolating the Cape Peninsula population. This temporal perspective is often missing from population genetic studies but is critical for understanding the evolutionary consequences of recent human activities. Our study reveals a striking contrast between contemporary and historical patterns of gene flow in caracal populations across our study areas. This temporal shift from connectivity to isolation provides compelling evidence for the genetic consequences of recent urbanization in this medium-sized carnivore.

### Historically connected populations

Few population genetic studies have been conducted on caracals, and to our knowledge, the only other studies in South Africa to use mitochondrial markers were conducted across a large geographic area in central and northeastern South Africa (Tensen et al. 2018, 2019). With their studies, Tensen and colleagues suggested that, based on mitochondrial markers, caracals across the entirety of South Africa are a single homogenous population. Our mitochondrial DNA findings support historically high levels of connectivity across large spatial scales in South Africa’s caracals, but the Cape Peninsula is now isolated by urbanization and not functionally part of the larger homogenous population.

The Cape Peninsula caracal population is characterized by several unique mtDNA haplotypes, but also has haplotypes shared with all three other study areas. These findings suggest the retention of historic signatures of connectivity both with caracals in the Greater Cape Town area, as well as caracal mtDNA lineages from further afield in the Central Karoo and Namaqualand as indicated by shared haplotypes between GCT, CK and NMQ. Network analysis of the haplotypes revealed genealogical relationships characteristic of extensive historic connectivity among caracal across the study region; identifying two clades within the dataset each comprising haplotypes from all four study areas and connected by haplotypes from NMQ and CK. This is further evidence for a pattern of regional-scale connectivity in caracals across South Africa (Tensen et al. 2019).

Patterns of haplotype and nucleotide diversity in the dataset indicate that all study regions support similar haplotype diversity. However, haplotypes within the Cape Peninsula and Namaqualand had the highest nucleotide diversity. In a multi-taxa study across China, mitochondrial DNA-based genetic diversity was negatively correlated with human population density (Hu et al. 2021), yet in the Cape Peninsula, nucleotide diversity currently remains high although it is surrounded by high human densities. Cape Peninsula nucleotide diversity is likely the result of how isolated CP caracals retain a combination of both unique haplotypes and those shared with GCT, CK and NMQ, as vestiges of historic connectivity across the study region. This pattern persists despite the strong signatures of contemporary genetic drift and differentiation observed in microsatellite and whole genome data across the four areas.

### Urban isolation, genetic drift and differentiation

Using microsatellite markers, the Cape Peninsula population exhibits clear signatures of genetic isolation, with significantly reduced allelic richness compared to our other study populations, relatedness values approximately double that of our comparative populations, and reduced expected heterozygosity compared with outgroup populations (GCT, CK, NMQ).

Collectively, these patterns are consistent with the effects of genetic drift in small, isolated populations following urbanization-induced habitat fragmentation (Delaney et al. 2010; Riley et al. 2014; Serieys et al. 2015; Miles et al. 2019; Schmidt et al. 2020; DeMarco et al. 2021). The retention of shared mtDNA haplotypes across all our study areas, including unique Cape Peninsula haplotypes, suggests that isolation occurred relatively recently, likely coinciding with the rapid urban expansion of Cape Town over the past several decades (Figure 2). A recent genomics study of the Cape Peninsula caracals also found evidence of strong genetic drift in the isolated population, estimating low levels of migration into the population over the past ∼75 years with increased isolation on the order of 33 years ago (Kyriazis et al. 2024). This finding is consistent with the progression of development in Cape Town (Figure 2). The genomics study also found long runs of homozygosity in the genomes of CP caracals, indicating reduced genetic diversity compared with the outgroups, further supporting that recent and rapid urbanization has had genetic consequences for CP caracals (Kyriazis et al. 2024). Genetic diversity and population differentiation have shown similar patterns in other isolated urban wildlife populations. In Germany, wild boar (*Sus scrofa*) exhibited distinct population differentiation among five urban forests compared with their rural counterparts (Stillfried et al. 2017). Mountain lions (*Puma concolor*) in southern California isolated by urban development and freeways also showed lower allelic richness and heterozygosity and increased relatedness among individuals (Riley et al. 2014). Genome-wide variation in the white-footed mouse (*Peromyscus leucopus*) was negatively correlated with the extent of urbanization, although the species thrives in urban environments (Munshi South et al. 2016). In a continent-wide North American study of the effects of urbanization on mammals and birds, mammals were found to consistently experience reduced genetic diversity compared with their counterparts living in wildland landscapes as a result the swift genetic drift that occurs when populations are isolated by urbanization (Schmidt et al. 2020). Thus, isolation by urbanization has, in numerous examples, led to a loss of genetic diversity as measured by multiple metrics in wildlife populations. This has implications for the adaptive potential of populations in a world with ongoing rapid environmental change (Exposito-Alonso et al. 2022).

The evidence for substantial recent genetic differentiation (using microsatellite loci) and population structuring, particularly between the rural populations and the isolated Cape Peninsula population aligns with predictions for small, isolated populations experiencing reduced gene flow and increased genetic drift. In a recent study in southern California (USA), the amount of impervious surface in the urban matrix was the greatest predictor of population structure and genetic differentiation across five bobcat populations (Kozakiewicz et al. 2019). In the same study areas, coyote populations also showed genetic structuring linked to urbanization.

Fragmented urban populations displayed clear differentiation, whereas no such structure was observed in connected non-urban areas, despite individuals in non-urban areas ranging over much larger distances than those confined to urban areas (Adducci et al. 2020). Although the

Cape Peninsula and the Greater Cape Town populations are in effect separated at their nearest points by approximately 3 km, the effect of the Cape Town urban matrix has reduced gene flow and caused sufficient isolation for a structuring effect. The barrier effect of the urban matrix and roads is clear from the few migrants detected in the Cape Peninsula population and from GPS-collar data (Supplemental Figure S1), which show that the dense Cape Town matrix restricts animal movements. This limited contemporary gene flow contrasts sharply with the historical connectivity indicated by shared mtDNA lineages.

The degree of genetic differentiation we observed is comparable to that reported for other carnivores in fragmented landscapes (Hellborg et al. 2002; Serieys et al. 2015; Basto et al. 2016) , suggesting that caracals respond predictably to habitat fragmentation despite their reputation for adaptability to diverse ecosystems. Of interest was that the greatest degree of genetic differentiation was between the Cape Peninsula population and the Central Karoo population, although the Namaqualand population is further from the Cape Peninsula than the Central Karoo population by approximately 200 km. Namaqualand is north of Cape Town and near a coastline while the Central Karoo and the Cape Peninsula and Greater Cape Town area straddle a significant mountain range, the Cape Fold mountains. These findings suggest that gene flow between Namaqualand and the Greater Cape Town region occurs more readily than between the Greater Cape Town and Cape Peninsula regions and the Central Karoo, possibly because of the elevated mountain range separating the Central Karoo and Cape Peninsula as opposed to low-lying, largely coastal areas separating Cape Town and Namaqualand. In a habitat selection studies in the Cape Peninsula, caracals were noted to select for low elevation areas and coastal areas (Leighton et al. 2022b; Serieys et al. 2023). Thus, barriers that include the Cape Fold Mountains and the Cape Town urban matrix impede gene flow and increase genetic differentiation.

The elevated relatedness values in the Cape Peninsula compared to other populations suggest that genetic drift has increased the frequency of related individuals within this isolated population. Specifically, it increases the shared allelic pool (which is getting smaller) such that individuals are more likely to share the same allele via both identity by descent (true relatedness) and identity by state. Importantly, while we observed reduced genetic diversity in the Cape Peninsula, inbreeding coefficients (FIS) did not differ significantly across populations, suggesting that inbreeding depression may not yet be a critical concern. However, continued isolation could lead to further genetic erosion and potentially inbreeding depression in future generations.

### Spatial ecology and relatedness

Our finding that individuals in the Cape Peninsula with higher pairwise relatedness also exhibit greater home range overlap provides important insights into the spatial organization of genetic structure within caracal populations. This positive correlation suggests that related individuals tend to have overlapping territories, which is consistent with patterns of natal philopatry or limited dispersal distances typical of solitary carnivores (Waser and Jones 1983). Yet the trend of home range overlap and relatedness was somewhat surprising and our findings were contrary to our hypothesis that relatedness would not influence the degree of home range overlap; within the Cape Peninsula, only 320 km^2^ of available habitat persists, and male home ranges in particular can be large (male average: 74 km^2^). The large home ranges combined with limited wildlife habitat availability has led to extensive overlap amongst male individuals. In contrast, only GPS-collared females (n = 3) in the southernmost section of our study area had overlapping home ranges, although we were unable to GPS collar every male and female in the Cape Peninsula. This spatial clustering of related individuals may accelerate local genetic drift and contribute to the development of fine-scale population structure, particularly in this fragmented habitat where dispersing individuals face an immense barrier to dispersal in the form of Cape Town’s urban matrix (Supplementary Figure S1). However, our observations of higher relatedness among those with overlapping home ranges is not unique, including among other carnivores. For example, related African wild dog (*Lyacon pictus*) packs had significantly greater peripheral overlap than non-related ones (Jackson et al. 2017). Female racoon (*Procyon lotor*) neighbors were more related than by chance alone (Ratnayeke et al. 2002). Also in the USA, mother-daughter bobcat pairs were found to have extensive home range overlap in a wildland-urban interface (Payne et al. 2024). We were unable to conduct additional analyses to test relationships among overlapping females because only three of our females overlapped home ranges. Nevertheless, the overall trend we detected is interesting considering that we did not observe dispersal out of the system while caracals were collared, although the structure plot suggests gene flow out of the Cape Peninsula into the GCT region.

## Conclusion

Our results contribute to the growing body of evidence that terrestrial mammals, and carnivores in particular, are highly sensitive to the genetic consequences of urbanization, demonstrating that urbanization can rapidly disrupt historical patterns of gene flow, leading to genetic isolation and erosion in isolated, urban-adjacent populations. The contrast between historical connectivity (indicated by shared mtDNA haplotypes) and contemporary isolation (revealed by microsatellite structure) underscores the recent and substantial impact of urban development on these caracal populations. The patterns we observed in Cape Town caracals mirror those documented in other urban carnivore populations demonstrating that physical barriers created by urban development can rapidly disrupt gene flow even between geographically proximate populations. With time, this population faces increased risk of inbreeding depression and reduced adaptive potential in the face of environmental change (Kyriazis et al. 2024) .

Our detection, albeit limited, of several possible first or second-generation migrants from Greater Cape Town into the Cape Peninsula indicates that some dispersal across the urban matrix is possible. This finding suggests that conservation strategies focused on enhancing connectivity, such as wildlife corridors, overpasses, or other landscape-scale interventions, could effectively restore gene flow and maintain genetic diversity in isolated urban populations. We suggest that the City of Cape Town prioritize the purchase of available land parcels and greenbelts (see Figure 2 for remaining undeveloped areas) within the urban matrix, and avoid further development of the False Bay coastline (Figure 2), to promote and maintain connectivity both in the short and longer term. Further, the City should actively manage those land parcels and greenbelts to protect wildlife remaining within the urban matrix from illegal poaching, which is an ongoing threat to wildlife in Cape Town.

Our study also contributes to the limited body of research examining the genetic impacts of urbanization on African wildlife populations. As urbanization continues to expand across Africa, understanding how native species respond to habitat fragmentation becomes increasingly critical for conservation planning. Our findings suggest that even highly adaptable, mobile species like caracals are not immune to both the ecological and evolutionary consequences of urban development, highlighting the importance of maintaining connectivity between urban and rural populations. The rural populations in our study serve as important genetic reservoirs, maintaining higher genetic diversity and potentially serving as sources for genetic rescue.

As global urban expansion continues to accelerate, three critical conservation strategies will be essential: 1) safeguarding sufficiently extensive and connected habitat fragments within urban areas to support viable wildlife populations, 2) reducing human-induced deterministic pressures such as human-induced mortality and habitat loss, and 3) establishing and sustaining ecological corridors within and among fragmented habitats to maintain metapopulation dynamics and ecosystem services in the urbanizing landscape (Crooks 2002).

## Author contributions

Laurel E.K. Serieys, Robert K. Wayne*, and Jacqueline M. Bishop contributed to the study conception and design. Sample collection was performed by Laurel E.K. Serieys, Marine Drouilly, Storme Viljoen, Bogdan Cristecu, Kristine J. Teichman, Deborah Winterton, and Gabriella Leighton. Laboratory analyses and data collection were performed by Shaelyn Squires, Megan Jackson, Laurel E.K. Serieys, and Jacqueline M. Bishop. Analyses were performed by Megan Jackson, Laurel E.K. Serieys, and Jacqueline M. Bishop. The first draft of this manuscript was written by Laurel E.K. Serieys and Jacqueline M. Bishop. All living authors commented on previous versions of the manuscript and all living authors read, edited, and approved the final manuscript.

*Robert K. Wayne, deceased December 26, 2022, was unable to comment on any version of the manuscript.

## Data availability

Mitochondrial sequence data will be made available through GenBank. Microsatellite data are available upon request.

## Acknowledgements

We are grateful for the logistical support of South Africa National Parks, the City of Cape Town, Cape Nature Conservation Board, and the Cape of Good Hope SPCA. We thank L. Mossop, J. Broadfield, C. Jansen, Dr. E Hellard and numerous volunteer interns for field assistance. Essential veterinarians were Drs. B. Stevens, A. Knight, E. Jordan, T. Hepburn. We also thank farmers of Namaqualand for sharing invaluable specimens and small-livestock farmers of the Central Karoo, including the Central Karoo Farmers Union, for allowing access to their farms and to carcasses of culled predators.

