## Supplemental Figure S1 for "Urbanization drives genetic erosion and population structure in a historically connected carnivore population"

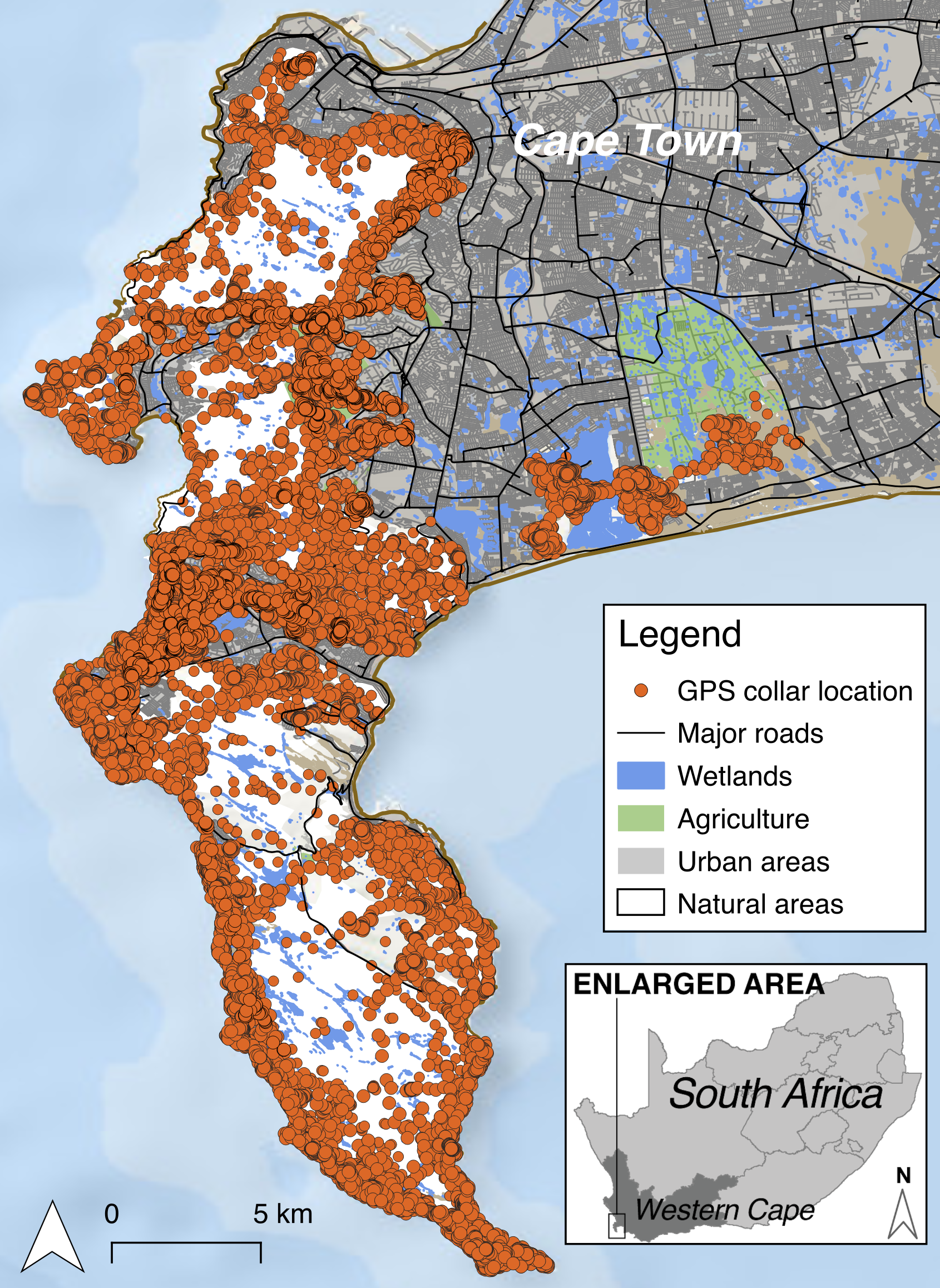
*Supplemental Figure S2. Cape Town’s urban matrix is a strong barrier to movement for GPS-collared caracals.*
